# Increased perceptual reliability reduces membrane potential variability in cortical neurons

**DOI:** 10.1101/2024.03.13.584630

**Authors:** Ben von Hünerbein, Jakob Jordan, Matthijs Oude Lohuis, Pietro Marchesi, Umberto Olcese, Cyriel M.A. Pennartz, Walter Senn, Mihai A. Petrovici

## Abstract

Uncertainty is omnipresent. While humans and other animals take uncertainty into account during decision making, it remains unclear how it is represented in cortex. To investigate the effect of stimulus reliability on uncertainty representation in cortical neurons, we analyzed single unit activity data recorded in mouse PPC, while animals performed a multisensory change detection task. We further used simulation-based inference (SBI) to infer membrane potential statistics underlying the spiking activity. Our analysis shows that stimulus changes increase spiking rate while decreasing its variability. The inferred membrane potential statistics suggest that PPC neurons decrease their membrane potential variability in response to task relevant stimuli. Furthermore, more perceptually reliable stimuli lead to a larger decrease in membrane potential variability than less reliable ones. These findings suggest that individual cortical neurons track uncertainty, providing Bayesian benefits for downstream computations.

## Introduction

The brain needs to make sense of the world based on noisy, unreliable, and ambiguous information. We experience this when having a conversation in a noisy room, finding our way in the dark or trying to pick our bicycle from a crowded rack. To reduce uncertainty we automatically incorporate information from additional sources; we look at people’s lips while they speak (Sumby and Pollack, 1954), use our hands to feel our environment in the darkness (Ernst and Banks, 2002), and use our memory to recall where we left the bike. To achieve these feats, our brain must represent and compute with the relative uncertainty associated with sensory percepts and prior knowledge.

This idea is often formalized using Bayesian principles, suggesting that the brain represents and computes information using probability distributions, for example, computing a posterior probability of a given event (Jaynes, 2003). Indeed a large body of work suggests that humans and animals take stimulus- and prior uncertainty into account when performing tasks such as coincidence detection (Miyazaki et al., 2005), gaze direction perception (Tassinari et al., 2006; Landy et al., 2012) and dynamic sensorimotor tasks (Faisal and Wolpert, 2009; Fetsch et al., 2009; O’Reilly et al., 2013; Meijer et al., 2019). Moreover, maladaptive uncertainty weighting of sensory and prior information has been linked to neurological disorders such as autism spectrum disorder and schizophrenia (Lawson et al., 2014; Van de Cruys et al., 2014; Goris et al., 2018; Stevenson et al., 2014; Noel et al., 2018). Understanding how the brain represents and computes with the uncertainty linked to different sensory perceptions is key to understanding cortical computation.

The posterior parietal cortex (PPC) has been identified as a hub for integrating information from different sensory modalities. Neurons in PPC have been found to respond to visual, auditory and somatosensory inputs (Raposo et al., 2012; Olcese et al., 2013; Mohan et al., 2018; Nikbakht et al., 2018). Indeed, the response of multisensory neurons to cross-modal stimulation is enhanced compared to the most effective of the individual stimuli (Stein and Stanford, 2008). Moreover, the strength of this “multisensory enhancement” depends crucially on several factors such as the temporal congruency (Meijer et al., 2017) and the perceptual strength of the individual stimulus components (Ibrahim et al., 2016). Such effects are expected from a Bayesian perspective: temporally proximal cues are likely to be informative of each other, and weaker cues leave more room for gaining additional information than strong cues. However, the mechanisms underlying this integration of information at the level of single neurons, and how information from different sources are weighted are still not well understood.

Single deep-layer cortical neurons receive bottom-up (Song et al., 2017) input driven by different sensory modalities and top-down input from higher order areas (Larkum, 2013; Rindner et al., 2022). They thus make an excellent substrate for combining novel sensory information with prior expectations. A recent theory (Jordan et al., 2021) offers a novel view on the dynamics of individual cortical neurons and suggests that they may be optimally suited to perform Bayes-optimal integration. This theory proposes that conductances play an important role in neuronal computation by allowing microscopic dendritic compartments to represent distributions with local, biophysical quantities. It suggests that neurons represent likelihood functions and priors in basal and apical dendrites respectively, while the soma computes the corresponding posterior distribution (Figure 1a). Specifically, the local effective reversal potential, the potential to which the membrane potential is pulled by synaptic inputs, represents the mean, while the sum of local excitatory and inhibitory conductances represent the precision of a Gaussian likelihood in each basal compartment. Intuitively, increasing synaptic conductances, e.g., by upstream stimulation, increase the pull towards the effective reversal potential. As this effect propagates to the soma it reduces sensitivity to unspecific background input. Computationally, this can be interpreted as sharper likelihood functions resulting in sharper posteriors. Biophysically, the variability of the somatic membrane potential is proportional to the posterior uncertainty. In line with a Bayesian view of brain function, the theory predicts that since new information reduces uncertainty, it also decreases membrane potential variability; furthermore, it predicts that the magnitude of this decrease is proportional to stimulus reliability. Finding evidence for such Bayesian processing at the single neuron level represents a promising step towards understanding how the brain deals with perceptual uncertainty.

**Figure 1:**
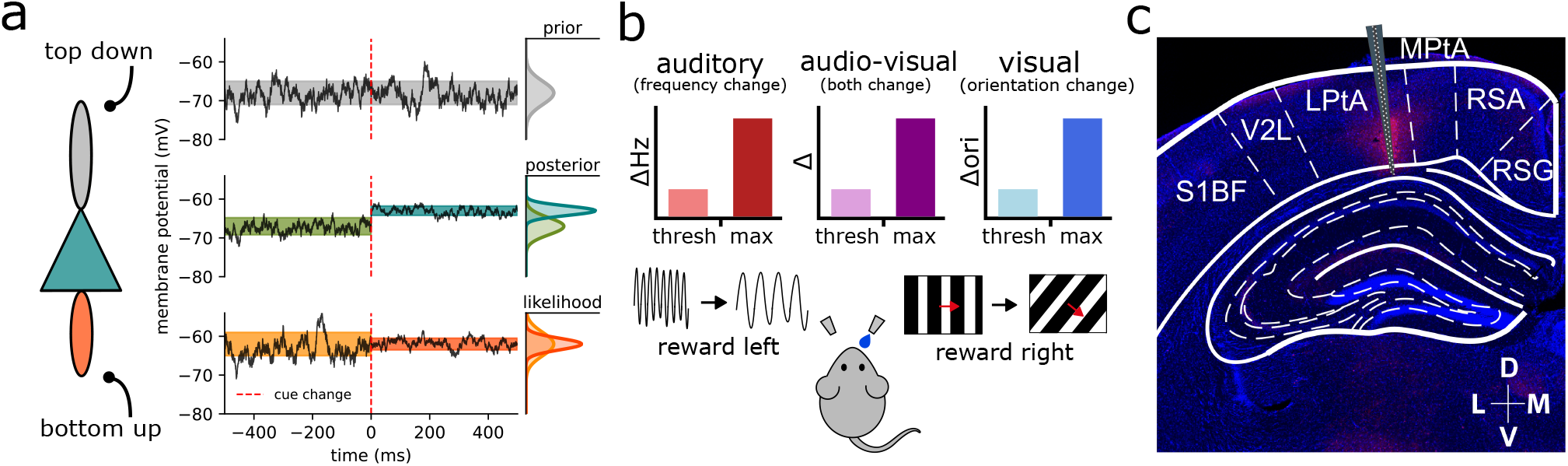
Perceptual reliability-weighted integration of sensory information in mouse PPC. (a) Conductance-based Bayes-optimal cue integration. According to Jordan et al. (2021), individual cortical neurons integrate novel (bottom-up) input with prior information (top-down) in a Bayes-optimal way at the level of somatic membrane potentials. Traces illustrate simulated membrane potentials of each compartment. The curves (right) represent the likelihood (basal dendrite, orange), prior (apical dendrite, grey), and posterior distribution (soma, green) pre- and post-stimulus. The presentation of novel information at 0 ms causes sharpening of the likelihood which reduces uncertainty in the posterior. (b) Behavioral task. A visual (drifting grating) and auditory (Shepard tone) stimulus were presented continously. Mice learned to report changes in either grating orientation (“V” trial, lick right), tone frequency (“A” trial, lick left) or both (“AV” trial, lick either side). Stimulus changes occurred either at maximum or threshold magnitude. (c) Single-unit activity was measured in mouse PPC. The image represents a reference section (Paxinos and Franklin, 2019) overlaid on the histologic verification with DAPI staining (blue) and electrode tract stained with DiI (red). LPtA, Lateral parietal association area; V2L lateral secondary visual cortex.

Here, we investigate neuronal responses in mouse PPC to uni- and multimodal changes of different magnitude in continuously presented auditory and visual stimuli (Figure 1b). First, we analyze how interspike interval statistics differ between pre and post stimulus change periods. Next, we infer and analyze membrane potential statistics underlying these spiking responses using simulation-based inference (SBI). Finally, we discuss our findings in the context of Bayesian computations in single neurons, and suggest future experiments.

## Methods

### Experimental Data

The data used in this study was collected to investigate the causal involvement of PPC in performing an audio-visual change detection task. For complete experimental details we refer to Oude Lohuis et al. (2022). All code used is openly accessible ^1^.

### Animals

Animal experiments were carried out in accordance with Dutch national legislation and institutional regulations. In total, 19 male mice were used from two transgenic mouse lines; PVcre (JAX 008069) and ai9-TdTomato cre-reported mice (JAX 007909). Mice were housed under a reversed day-night schedule (lights were on from 20:00PM until 8:00AM). As mice are nocturnal, experiments were performed during the dark period. Animals were aged at least 8 weeks at the beginning of the experiments.

### Head bar implantation

Animals were implanted with a head bar prior to experimentation. Animals were anesthetized using isoflurane (3% for induction, 1.5 − 2% for maintenance) and fixed in a stereotaxic apparatus. After exposing the animals skull, a circular titanium head bar was glued and cemented to it. Mice were given 2 − 7 days to recover following surgery. Prior to the training procedure mice were habituated to the head fixation and the handling of the experimenter. A subset of animals underwent additional surgery to virally infect individual areas for opsin expression for optogenetic manipulations. However, data collected during sessions with optogenetic manipulations were not used in this study.

### Audiovisual change detection task

Mice were deprived of water and earned their daily ration by performing the behavioral task. Lick spouts were positioned within tongue reach symmetrically on the left and right side of the head-fixed animals. Auditory and visual stimuli were presented continuously throughout each behavioral session (Figure 1b).

The visual stimulus consisted of a drifting square-wave grating with a temporal frequency of 1.5 Hz and a spatial frequency of 0.08 cycles per degree at 70% contrast. The auditory stimulus was a stationary Shepard tone (Shepard, 1964), consisting of a center tone combined with its two lower and higher harmonics. Center tones used throughout the experiment ranged a one octave from 2^13^ Hz (8, 372 Hz) to 2^14^ Hz (16, 744 Hz). The weight of the center and harmonic tones were taken from a Gaussian distribution over weights, here centered at 2^13.5^ Hz (11, 585 Hz). Tones were high-pass filtered (Beyma F100, Crossover frequency of 5 − 7 kHz) and presented at a sampling rate of 192 kHz using two bullet tweeters (300 W) positioned directly below the screen. The sound pressure level was calibrated at the position of the mouse, and the volume was adjusted per mouse to the minimum volume that maximized performance (on average around 70 dB).

In visual change trials (“V trials”), the orientation of the drifting grating was instantaneously changed while preserving the phase. Likewise, in auditory change trials (“A trials”) the stimulus was instantaneously changed from one Shepard tone to another, with different center frequency and associated harmonics. Animals were trained to respond to changes in a lateralised manner (A trial: lick left, V trial: lick right). Modality-side pairing was counterbalanced across mice. Thus, to successfully perform the task, the animal had to monitor both auditory and visual modalities to detect changes in either.

During training, trial types were ordered pseudo-randomly by block-shuffling; consecutive blocks of 10 trials each were comprised of randomly ordered trial types in a fixed proportion (8% catch, 46% V trials, 46% A trials). During testing, audio-visual change trials (“AV trials”) were introduced, where both auditory and visual stimuli changed (8% catch, 33.5% A trials, 33.5% V trials, 25% AV trials). AV trials were rewarded for the first lick on either side. Thus, in contrast to A and V trials, cues in AV trials were not informative about the response they should elicit.

Additionally, to manipulate their perceptual reliability, changes in either of the two modalities were delivered either at perceptual threshold or at maximum magnitude. The perceptual threshold for each modality was determined for each animal separately by fitting a psychometric function over the accuracy achieved across five levels of stimulus change magnitude. In trials with an auditory change, the center pitch changed by 1/10^th^ of an octave in threshold and a full octave on in maximum change trials. In trials with a visual change the movement orientation changed by 4 − 7 degrees in threshold and 90 degrees in maximum change trials. Consecutive trials were separated by inter-trial intervals sampled randomly from an exponential distribution with a mean of 6 s, a minimum of 2 s and maximum of 20 s. Following a stimulus change, mice were rewarded if the first lick after 100 ms and before 1500 ms was made to the correct side. Sessions were terminated after 20 trials of unresponsiveness, with these last 20 trials being discarded from all analyses. Mice achieved a median reaction time of 324 ms across auditory and 407 ms across visual hits.

### Neural recordings

Mice were anesthetized (isoflurane, induction at 3%, maintenance at 1.5–2%) and small craniotomies (approx. 300 − 500 *μ*m) over the areas of interest were made using a dental drill. Areas of interest were identified based on stereotaxic coordinates (V1: AP 0.0, ML *±*3.0, PPC: AP 1.9, ML *±*1.6, AC: AP 2.6, ML *±*4.3, relative to lambda and bregma, respectively) (Goard et al., 2016; Song et al., 2017; Le Merre et al., 2018). Extracellular recordings were performed on consecutive days with a maximum of four days to minimize cortical tissue damage. Microelectrode arrays (Figure 1c) with 32 or 64 channels (NeuroNexus, A1×32-Poly2-10 mm-50s-177, A1×64-Poly2-6 mm-23s-160) were slowly inserted either perpendicularly to the cortical surface (V1), or at an angle of approximately 30 degrees away from the midline (AC) After insertion, the probe was left in place for at least 15 min before recording to allow the tissue to stabilize. To identify the exact probe location post hoc, the probe was covered in Dil (Thermo Fisher Scientific) on the final day of recording. Neurophysiological signals were pre-amplified, bandpass-filtered (0.1 Hz to 9 kHz) and acquired continuously at 32 kHz using a Digital Lynx 128-channel system (Neuralynx). Post experiment, mice received an overdose of pentobarbital, were transcardially perfused (4% PFA in PBS) and their brains were extracted.

### Neurophysiological data processing

Spike sorting was performed using Klusta software and manually curated using the Phy GUI (Rossant et al., 2016). Prior to spike sorting, common noise artifacts were eliminated by subtracting the median of the raw trace of surrounding channels (within 400 mm). Each potential single unit was inspected based on its waveform, autocorrelation function, and firing pattern across channels and time. Single units were only included if they met the following criteria: (1) isolation distance *>* 10 (Schmitzer-Torbert et al., 2005); (2) 0.1% of their spikes occurred inside the 1.5 ms refractory period (Vinck et al., 2016; Bos et al., 2017); and (3) stable presence throughout the session. To asses stability the session-wise activity was binned and single units were considered stable if they fired in at least 90% of bins.

### Pre-processing

We split the trial-wise activity into a pre- and post-stimulus period. The pre-stimulus period spanned the 1500 ms before the stimulus change, while the post-stimulus period covered 200 ms to 1700 ms after the change. Due to transient responses in the initial 200 ms post-stimulus, we excluded this interval to better capture sustained within-trial statistics following stimulus onset.

Accordingly, we summarised the trial-wise spiking activity in each period by computing the mean and standard deviation of the inter-spike interval statistics (ISI, *μ*_s_, *σ*_s_). Note that this summary statistic assumes that the ISIs are unimodally distributed and thus would not adequately describe bimodal distributions. We thus quantified the amount of bursting within each spike train (Chen et al., 2009) and removed spike trains with excessive bursting (see also Supplementary Figure 2). Finally, as the inference procedure is less accurate for spike trains with fewer spikes (see also section Supplementary Figure 3), we only included trials with at least 10 Hz in both pre- and post-stimulus change periods in our analysis. Note that this may bias the sample towards neuron types associated with higher firing frequencies, namely interneurons and deep cortical pyramidal neurons (Contreras, 2004). After filtering, 4170 trials collected from 83 neurons in 6 animals remained for subsequent analysis (*V* = 1674, *A* = 1663, *AV* = 524, catch = 309; non-catch trials balanced across max and thresh change magnitudes).

## Simulation-Based Inference (SBI)

SBI can be used to efficiently compute an approximate posterior distribution in cases where the likelihood function is intractable, e.g., stochastic simulations. This is achieved by leveraging universal function approximators to represent distributions (Tejero-Cantero et al., 2020). A typical application of SBI proceeds as follows: First, we choose an appropriate mechanistic model and a prior distribution over its parameters (see Figure 2). Next, we simulate the model with random samples from the prior distribution and record observations. We then train a density estimator to map recorded observations to the underlying parameters. Finally, we can use our trained density estimator to compute approximate posteriors over parameters for new observations, in our case spiking data obtained from experiments. We used SBI to infer membrane potential statistics of biological neurons by inverting the dynamics of leaky-integrate and fire (LIF) neurons using Sequential Neural Posterior estimation (SNPE; Greenberg et al., 2019). We implemented the SBI workflow using the sbi toolbox^2^ in Python.

**Figure 2:**
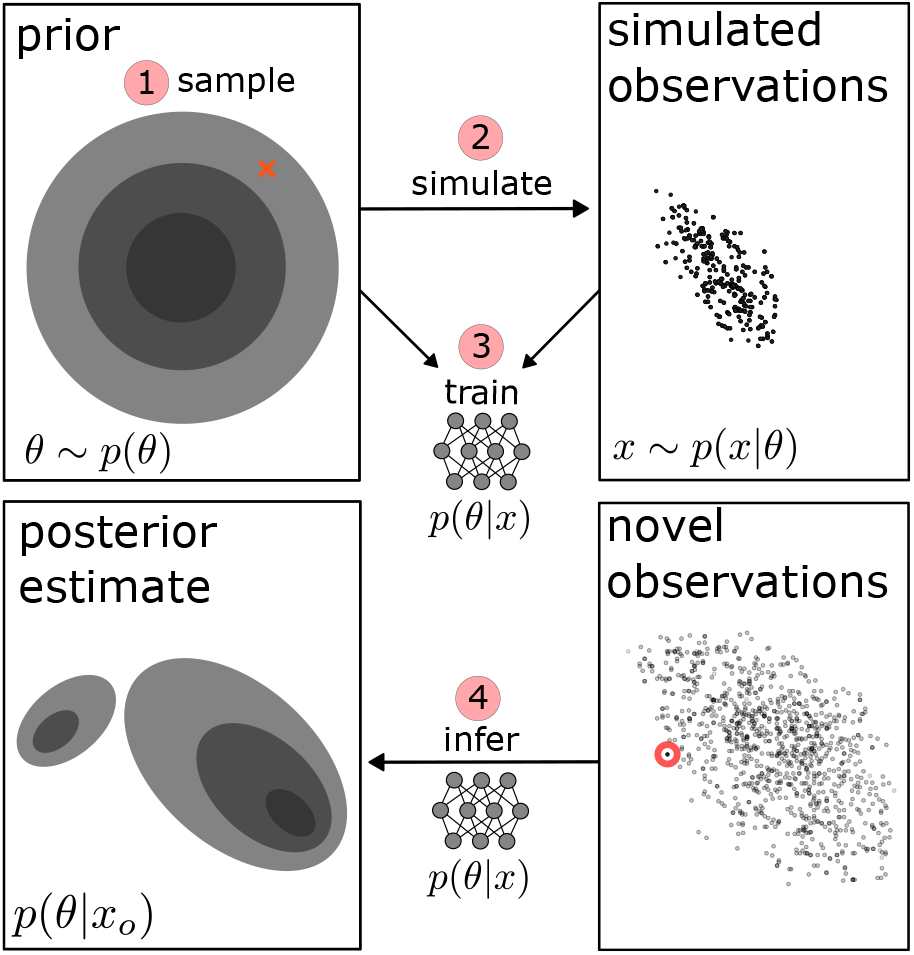
Simulation-based inference (SBI). SBI is a method for recovering the hidden parameters underlying novel observations. (1) A single sample (red cross) from the prior distribution over the parameters (*θ*) of the mechanistic model. (2) Using this sample, observations *x* are generated by (stochastic) simulation. (3) The parameters and observation pairs are then used to train a density estimator to learn an approximate posterior distribution *p*(*θ*|*x*), i.e., to invert the simulation probabilistically. (4) The trained density estimator is used to infer a posterior distribution over the underlying parameters of a novel observation (*x*_o_). Here, SBI was used to infer the parameters *θ* = (*μ*_I_, *σ*_I_) of the input current *I* underlying the observation generated from a single spike train 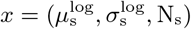.

### Mechanistic model

We chose the leaky-integrate-and-fire (LIF) neuron model due to its simplicity, ease of simulation, and common use in neuron modelling. Its dynamics is given by

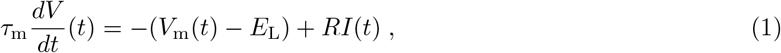

with the additional threshold condition

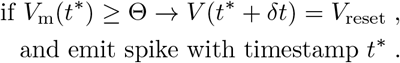

Here, *V*_m_ is the neuron’s membrane potential, *τ*_m_ the membrane time constant, *E*_L_ the reversal potential of the leak current, *R* the membrane resistance, Θ the spiking threshold and *V*_reset_ the reset potential. Due to the spiking mechanism a hard threshold on the membrane potential with subsequent reset, the membrane potential statistics of a LIF neuron are difficult to parameterize directly (Petrovici et al., 2016). Here we instead chose to parameterize an input current with Gaussian noise

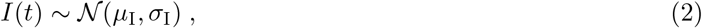

with mean *μ*_I_ and standard deviation *σ*_I_. This current subsumes all synaptic inputs to the neuron. All network simulations were carried out with NEST^3^ (Gewaltig and Diesmann, 2007; de Schepper et al., 2022).

### Parameter prior

The prior defines the space of parameters used to simulate the mechanistic model. It thus implicitly defines the support (bounds) of the posterior distribution. We initialized the prior as a uniform distribution over the range [−2 pA, 2 pA] for the mean *μ*_I_ of the injected Gaussian noise current, and [0 pA, 30 pA] for its standard deviation *σ*_I_. The prior was then iteratively restricted by removing parts of the parameter space that yielded no or excessive firing. Here we define “excessive firing” as firing rates larger than the maximum observed in the data.

### Observables

Spike trains were the common quantity produced by simulations and experiments. Here, we summarized an individual spike train using the empirical mean, standard deviation and raw number of spikes 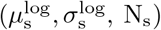 of the log inter-spike interval (ISI) distribution (Figure 3a,b). We chose a summary statistic because we did not want to infer parameters that reproduce the exact spike times but only the spiking statistics. We log transform ISIs because it reduces the posterior uncertainty of our inference procedure.

**Figure 3:**
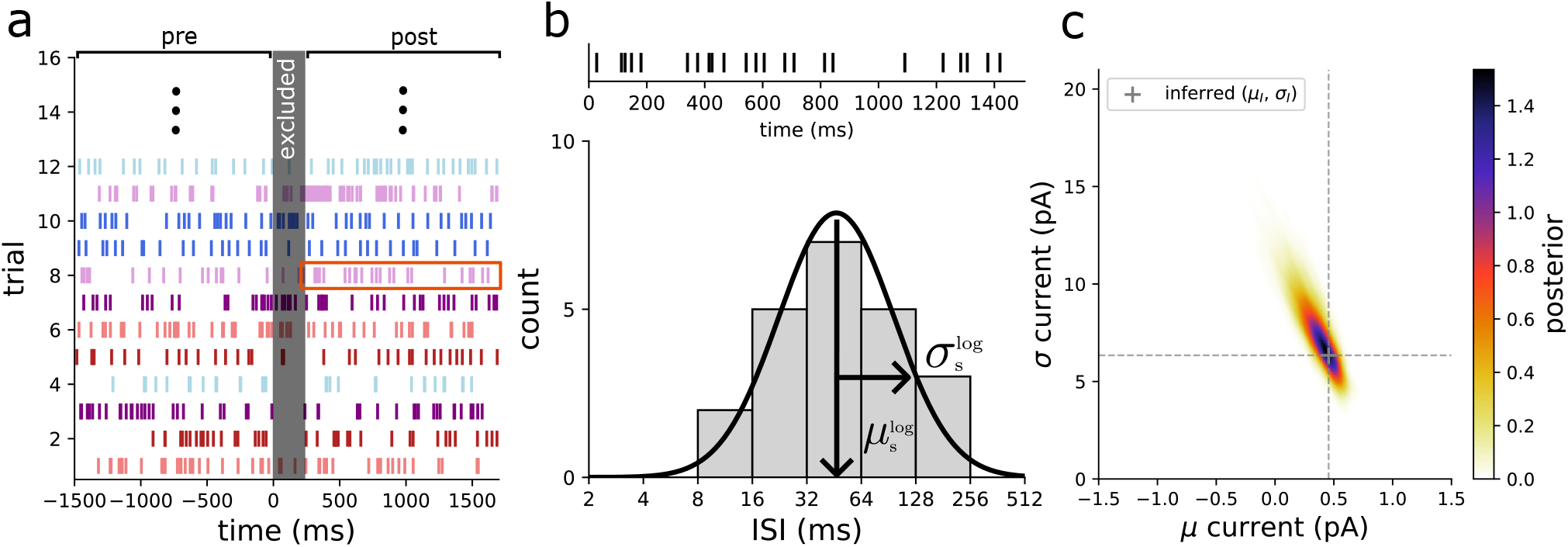
Inferring intracellular quantities from spike trains. (a) Single PPC neuron spiking activity recorded during successive trials separated into pre- and post-stimulus change spiking activity. Colors denote trials from different conditions (see Figure 1). (b) Inter-spike interval (ISI) statistics. Each spike train is summarized by the mean and standard deviation of its log ISI distribution. (c) Inferring trans-membrane currents. The density estimator trained with SBI is used to infer an approximate posterior distribution over input current parameters from each observation (*μ*_s_, *σ*_s_). For subsequent analysis we collapse this distribution to its most likely values (grey cross).

### Training

We trained a density estimator (Lueckmann et al., 2019; Papamakarios et al., 2019) to learn the mapping from observations to current parameters. To generate training data, i.e., parameter/observation pairs, we sampled parameters from the restricted prior distribution, performed a simulation of the neuron model and computed the respective observation from the simulation output. We simulated each neuron for 2000 ms, the first 500 ms of which were discarded to allow the neuron to reach its steady state. The spiking activity in the remaining 1500 ms was recorded. We generated and trained the mixed density estimator on 150, 000 parameter/observable pairs.

### Validation

To validate our learned density estimator, we sampled 20, 000 parameter pairs, not seen during training, from the prior. We simulated each parameter pair and recorded the observations. Next we performed the inference procedure for each simulated observation and computed the difference between the inferred and the original parameters. To determine the influence of the number of spikes on inference quality, we computed the errors separately for varying spike count. We found that the inference procedure was biased towards underestimating *μ*_I_ while overestimating *σ*_I_ at firing rates below 10 Hz (see also Supplementary Figure 3). This lower boundary of 10 Hz was used as an exclusion criterion for recorded trials.

### Inference

To infer the parameters underlying single recorded spike trains we conditioned the posterior on a single observation. We collapse this distribution to its most likely value to obtain the parameters most likely underlying the observation (Figure 3c). In a final step we translated these current parameters to membrane potential statistics by simulating a neuron with the inferred parameters (*μ*_I_, *σ*_I_) and recording the membrane potential traces from which we can compute the relevant statistics (*μ*_u_, *σ*_u_, where u indicates that this is a quantity relating to membrane potentials).

## Statistical Analysis

We used linear mixed modelling (LMM) in SPSS 29.0 to investigate the effect of change magnitude and modality both on the ISI and the inferred voltage statistics. We chose LMMs because they are robust to imbalanced designs and are able to account for the nested structure of the data (Aarts et al., 2014). LMMs describe the relationship between a response variable and multiple explanatory variables of two types; fixed effects are the independent variables (change magnitude, modality) whereas random effects are grouping variables that account for the nested structure of the data. Here, the nested structure arose due to repeated trials across conditions from the same animals. In our statistical model, we defined modality and change magnitude as our fixed effects and included animal identity as our random effect. We chose a covariance structure whereby the intercept for each animal was allowed to vary freely. We did not include the neuron identity from which trials were recorded as a nested random effect within animals as the sometimes limited number of trials recorded per neuron (*>* 5) meant intercepts could not be estimated reliably and the model would not converge. For our dependent measure, we computed the change both for the recorded ISI statistics 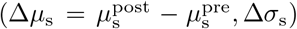 and the inferred membrane potential statistics (Δ*μ*_u_, Δ*σ*_u_). The following statistical model was fitted for each of these dependent measures *y ∈ {*Δ*μ*_s_, Δ*σ*_s_, Δ*μ*_u_, Δ*σ*_u_*}*:

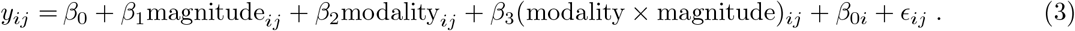

Here *y*_*ij*_ is the *j*^*th*^ observation within the *i*^*th*^ animal. The overall intercept (bias) is denoted by *β*_0_. The coefficients *β*_1_, *β*_2_ and *β*_3_ correspond to the coefficients for magnitude, change modality and their interaction respectively. *β*_0*i*_ denotes the random intercept for each animal and *E*_*ij*_ represents the residual error. For main effects we report the F-test as F_(*df*1,*df*2)_ = F where *df* 1 and *df* 2 indicate the between- and within-group degrees of freedom respectively.

## Results

### ISI Statistics

We first analysed the effects of stimulus change and magnitude on the ISI statistics recorded during pre- and post-change period in mouse PPC (Figure 4a,b). Here, we expected stimulus changes to results in a decrease in within-trial spiking variability. Indeed, we found that across all conditions stimulus change led to a decrease in the mean inter-spike interval (F_(1,7.4)_ = 31.72; *p <* 0.001) and a decrease in the standard deviation of the inter-spike interval (F_(1,3.9)_ = 23.92; *p* = 0.008). Further, we expected that larger change magnitudes would have a stronger effect on neural activity. While maximum change trials appeared to have a lower *μ*_s_ than threshold trials, this difference was not significant. However, max change trials showed a stronger decrease in *σ*_s_ than threshold change trials.

**Figure 4:**
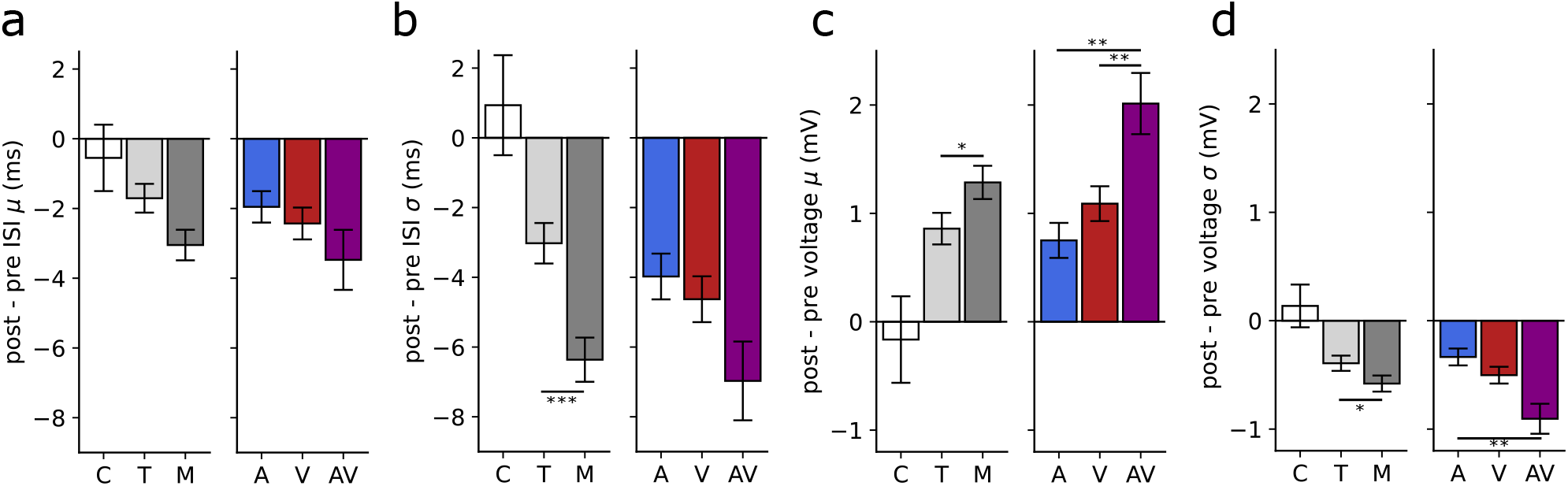
Effect of stimulus change on within trial ISI and inferred membrane potential statistics. (a) The difference in post- and pre-stimulus mean (*μ*_s_) and (b) standard deviation (*σ*_s_) of the recorded inter-spike interval distributions across change modalities and for (T)hreshold and (M)ax change magnitudes. (c) The difference in post- and pre-membrane potential mean and (d) standard deviation (right) as inferred by SBI. Error bars indicates SEM and *, ** and *** indicate significance at the 0.05, 0.01 and 0.001 levels respectively.

Further, we expected a difference between uni- and multi-modal stimulus changes as unimodal stimuli were matched for intensity. While a trend in this direction could be observed, AV trials did not differ significantly from unimodal A and V trials on the change in the mean or standard deviation of the ISI statistics. One reason for this could be that we are lacking statistical power due to the smaller number of AV trials compared to A and V trials. In correct rejection (catch) trials we did not observe a significant change in the ISI statistics between pre and post change (one sided t-test; Δ*μ*_s_, *p* = 0.282, Δ*σ*_s_ *p* = 0.257,).

**Table 1:** Effect of stimulus change on ISI statistics.

|  | Conditions<br>(a vs b) | Mean difference<br>(a - b) | Standard<br>Error | Significance | CI<sub>95%</sub><br>[lower, upper] |
| --- | --- | --- | --- | --- | --- |
| $\Delta\mu_s$ | Max vs Thresh | -0.830 | 0.695 | 0.233 | [-2.193, -0.533] |
|  | A vs V | 0.398 | 0.647 | 1.000 | [-1.151, 1.848] |
|  | AV vs A | -1.312 | 0.945 | 0.496 | [-3.576, 0.952] |
|  | AV vs V | -0.913 | 0.944 | 1.000 | [-3.174, 1.347] |
| $\Delta\sigma_s$ | Max vs Thresh | -3.196*** | 0.991 | 0.001 | [-5.140, -1.253] |
|  | A vs V | 0.502 | 0.922 | 1.000 | [-1.708, 2.707] |
|  | AV vs A | -2.914 | 1.354 | 0.094 | [-6.156, 0.328] |
|  | AV vs V | -2.412 | 1.351 | 0.223 | [-5.648, 0.824] |

### Membrane potential statistics

Next, we analysed the membrane potential statistics underlying these changes in spiking statistics as inferred using SBI (Figure 4c,d). We expected, in line with the theoretical prediction, that a stimulus change would be associated with a decrease in the membrane potential variability. Indeed we found that stimulus change led to an increase in the mean membrane potential (i.e., a depolarization) (F_(1,4.3)_ = 30.804, *p* = 0.003) and a decrease in the membrane potential variability (F_(1,5.2)_ = 30.798, *p* = 0.002). Furthermore, we found that this effect of stimulus change associated increases in *μ*_*u*_ and decreases in *σ*_*u*_ was larger in maximum than in threshold change trials. Finally, we analysed the effect of change modality on membrane potential statistics. The comparison of uni- and multimodal change trial effects shows that AV trials led to a larger mean depolarization than A but not V trials.

That the above observed increase in firing (decrease in *μ*_s_) is associated with a decrease in *σ*_u_ is unexpected in light of our mechanistic neuron model. This is because in the fluctuation driven regime (Kuhn et al., 2004) larger *σ*_u_ will make it more likely for the membrane potential to reach threshold, leading to an increase in firing. However, this finding aligns with the prediction that integration of information leads to decreases in the membrane potential variability. These results support the predictions from Jordan et al. (2021) whereby individual neurons respond to more informative stimuli, i.e., larger cross-modal changes, by reducing their membrane potential variability.

**Table 2:** Effect of stimulus change on membrane potential statistics.

|  | Conditions<br>(a vs b) | Mean difference<br>(a - b) | Standard<br>Error | Significance | CI<sub>95%</sub><br>[lower, upper] |
| --- | --- | --- | --- | --- | --- |
| $\Delta\mu_u$ | Max vs Thresh | 0.527* | 0.244 | 0.031 | [0.049, 1.006] |
|  | A vs V | -0.315 | 0.227 | 0.495 | [-0.859, 0.229] |
|  | AV vs A | 1.145** | 0.333 | 0.002 | [0.348, 1.943] |
|  | AV vs V | 0.830** | 0.332 | 0.007 | [0.034, 1.626] |
| $\Delta\sigma_u$ | Max vs Thresh | -0.240* | 0.118 | 0.042 | [-0.471, -0.008] |
|  | A vs V | 0.098 | 0.89 | 0.810 | [-0.115, 0.312] |
|  | AV vs A | -0.520** | 0.161 | 0.004 | [-0.905, -0.134] |
|  | AV vs V | -0.363 | 0.161 | 0.072 | [-0.747, 0.022] |

## Discussion

### Summary & interpretation

Here we investigated the effect of stimulus-associated reliability on the within-trial inter-spike interval and membrane potential statistics of single neurons in mouse PPC. The stimulus change associated reliability was manipulated (low vs. high) by adapting the magnitude (thresh vs. max) and modality (A, V vs. AV). We found that stimulus changes across conditions led to increased and more regular spiking. Further, our results demonstrate that the reduction in ISI variability was proportional to stimulus reliability, i.e., larger (max) stimulus changes led to a stronger reduction. However, we did not observe a multisensory enhancement (Stein and Stanford, 2008) at the level of spiking statistics. Apart from a lack of statistical power due to the imbalance in single and multimodal trials, this may be due to the stimulus components in AV trials not being mutually informative. As animals learned to respond to unimodal change stimuli in a lateralized way, multimodal stimuli may have led to ambiguity with respect to the desired response (see Methods). This stands in contrast to previous studies showing multi-modal enhancement where the integration of cross-modal stimuli was beneficial for task performance (Olcese et al., 2013; Nikbakht et al., 2018; Meijer et al., 2018). On the other hand, at the level of inferred membrane potential statistics, stimulus changes across all conditions lead to an increase in the mean and a decrease in the standard deviation of the membrane potential distribution. Moreover, this effect appeared to be sensitive to stimulus reliability as it was stronger in multi-modal and maximum change magnitude trials compared to unimodal and threshold magnitude changes respectively.

Our results suggest that neurons in PPC are sensitive to the stimulus associated perceptual uncertainty at the level of their membrane potential statistics. Previous work found a whisking-induced reduction of membrane potential variability in rat primary sensory cortex (Crochet et al., 2011; Yamashita et al., 2013; Poulet and Crochet, 2019). Although they did not investigate the effect of reliability, these findings are in line with ours, suggesting that this effect is a widespread cortical phenomenon. Secondly, we show that the effect is sensitive to the stimulus associated reliability, suggesting that individual neurons track the stimulus related uncertainty. In light of recent theoretical work (Jordan et al., 2021), this stimulus-driven reduction can be viewed as a signature of Bayesian computation at the single-neuron level. In a Bayesian framework, additional information leads to a reduction of uncertainty, explaining the post-change decrease of variability. Furthermore, more informative (e.g., more reliable or salient) stimuli lead to a larger reduction of uncertainty and thus to a larger decrease of membrane potential variability. In contrast to previous work, we thus suggest that the stimulus-driven reduction in membrane potential variability is a consequence of single-neuron computation, rather than an emergent property at the network level (Sussillo and Abbott, 2009; Deco and Hugues, 2012). Note that here we focused on within-trial variability, rather than trial-by-trial variability. While their relationship is complex, a reduction of within-trial variability can cause a reduction in trial-to-trial variability. We suggest that the well-documented reduction in stimulus-driven trial-by-trial variability (Finn et al., 2007; Churchland et al., 2010; Wright et al., 2017) could in part be attributed to the within-trial activity reductions observed here.

### Probing the theory further

In the theory proposed by (Jordan et al., 2021), the stimulus-associated reliability is represented via local membrane conductances. Future work could apply SBI to a more complex neuron model including transmembrane conductances in addition to transmembrane currents. This would account for neuronal morphology by more accurately covering the effect of both distal (effectively current-based) and proximal (effectively conductance-based) synapses on the somatic membrane. Explicitly inferring a change in total input conductance following a change in the stimulus would lend further support to the concept of Bayesian processing at the single neuron level.

Furthermore, one could separately investigate the reliability-modulated contributions of presynaptic activity and synaptic coupling strength to the synaptic conductances. Whereas changes in the presynaptic rate lead to quick re-weighting across modalities (e.g., Fetsch et al., 2012), changes to the synaptic coupling reflect more long-term changes. These could be triggered by changes in reward probabilities or changes in sensory-sensory associations (Knöpfel et al., 2019). Future experiments could disentangle these components by using SBI together with a model where they are parameterized separately.

Alternatively, inferring intracellular parameters across time, rather than only their time-averaged statistics as we have done here, may allow for distinguishing between different models of biophysical computation. For example, probabilistic population codes (PPCs; Ma et al., 2006) consider a feedforward integration at the population level in which the respective efferent synaptic weights determine the relative weight of populations representing information from different modalities. Similarly, feedback normalization models (Ohshiro et al., 2011) assign effective “dominance weights” to afferents of multisensory neurons. These models thus exclusively rely on long-term plasticity to adapt to changes in relative reliability between modalities. In contrast, the Bayesian-dendrite theory considered here represents reliability via conductances. These are influenced by both synaptic weights and presynaptic rates and thus allow for rapid adaptation via presynaptic modulation of firing rates. Investigating how closely the neurally represented reliability follows rapid changes in stimulus-related reliability would allow us to differentiate between these model classes.

### Proxies for reliability

In this study we manipulated the perceptual reliability of the stimuli by varying the magnitude of the change. The task-relevant cue consisted of a change in one of two continuously presented stimuli. The reason we chose stimulus change as our cue (as opposed to stimulus onset) was to manipulate the stimulus reliability separately from its perceptual intensity. Thus, while larger stimulus changes were more reliably informative towards the availability of reward, the perceptual intensity, i.e., sound volume or visual contrast, of the stimuli did not change. However, given that neurons adapt to continuous stimulation (Kohn, 2007; Ulanovsky et al., 2003) it may be that stimulus change was associated with a perceived change in intensity. It would be interesting to explore whether the effects observed here are maintained under experimental manipulations where the stimulus reliability is fully decoupled from, or even inversely related to, perceptual intensity. Based on Bayesian principles we would expect that, independent of the way reliability is modulated, increasing reliability should decrease membrane potential variability. Here we only probed two levels of change magnitude per modality. It would be interesting to investigate whether neuronal responses are sensitive to more finely graded reliabilities as Bayesian principles would suggest a monotonic decrease of variability with reliability.

### Validation with ground-truth data

Here we used simulation-based inference together with simulations of a LIF model parameterized by a Gaussian current input to obtain single neuron membrane potential statistics underlying spiking activity. We validated our approach in simulation by inferring known parameters, both to assess inference quality and determine the amount of information needed to correctly infer model parameters. To further strengthen the inference procedure one would ideally compare the quantities inferred from recordings to a ground truth. This was not feasible in the current work as it would require, simultaneously recorded intra- and extracellular data which are challenging to obtain, especially over longer periods. In addition to providing a ground truth, these parameter and observation pairs recorded in vivo could be used to fine tune the density estimator used for the inference procedure using transfer learning (e.g., Finn et al., 2017). Establishing such ground truth benchmarks would allow researchers to test model predictions about computationally relevant but difficult to access quantities in a standardized way.

### Leveraging SBI to test theoretical predictions

Computational theories are often formulated with normative principles and mathematical convenience in mind. Thus many theoretical predictions pertain to abstract and difficult-to-access quantities. Testing these predictions in experiments is often not feasible due to cost and time constraints. As demonstrated here, SBI provides a way to bridge this gap by fitting the relevant quantities in mechanistic neuron models (Gonçalves et al., 2020). These inferred parameters can then be interpreted to gain valuable insights about the mechanisms underlying otherwise obscure phenomena. SBI could be particularly useful for testing the implicit predictions of microcircuit models which describe cortical computation as the product of canonically arranged groups of neurons (Sacramento et al., 2018; Wilmes and Clopath, 2019; Hertäg and Clopath, 2022; Haider et al., 2021; Granier et al., 2023).

With SBI, these predictions could be more readily tested based on existing data. For example, a recent model for how cortical microcircuits may implement efficient learning by approximating the backpropagation of errors algorithm (Sacramento et al., 2018) suggests that apical compartments receive lateral and top-down information to compute local errors (Fişek et al., 2023). This prediction is challenging to test as it requires in vivo recordings from apical dendrites. However, synaptic inputs to cortical pyramidal cells form a considerable contribution to local field potentials (LFPs) (Einevoll et al., 2013). One could use SBI together with existing simulation tools for microcircuits and their associated LFPs (Skaar et al., 2020; Rimehaug et al., 2023) to infer net currents flowing into apical dendrites during task learning. If apical compartments represent errors, these currents should shrink over the course of learning. Likewise one could test the prediction by more recent work (Haider et al., 2021) that apical and somatic compartments receive strongly correlated inputs.

## Conclusion

In this work we investigated how cortical neurons respond to uni- and multi-modal stimuli associated with varying perceptual reliabilities. We demonstrated that task relevant information decreases variability both on the spiking as well as the inferred membrane potential level. As predicted by a recent Bayesian model of single neuron computation we found a reduction in variability in proportion to the stimulus associated reliability. Our results shine new light on the representation of uncertainty in cortex and suggest that individual cortical neurons keep track and compute with uncertainty.

## Acknowledgements

This work has received funding from the European Union 7th Framework Programme under grant agreement 604102 (HBP), the Horizon 2020 Framework Programme under grant agreements 720270, 785907 and 945539 (HBP), the Swiss National Science Foundation (Sinergia grant CRSII5-180316), the Manfred Stärk Foundation, FLAG-ERA JTC projects CANON (2015) and DOMINO (2019), both cofinanced by the Netherlands Organization for Scientific Research. We also thank Sietse Kiewiet for helpful contributions to the manuscript.

## Footnotes

1 https://github.com/unibe-cns/reliable-membranes

2 www.mackelab.org/sbi/

3 www.nest-simulator.org

