## Supplement Information for "Increased perceptual reliability reduces membrane potential variability in cortical neurons"

March 13, 2024

### 1 Bursting

We computed a bursting metric based on (Chen et al., 2009) for each pre- and post-stimulus spike train. Here each spike train is given by given by,

$$a(t) = \sum_{n=1}^{N} \delta(t - t_n), \tag{1}$$

With $t_1, \cdots, t_N$ denoting the series of spike times, and N being the number of spikes. The measure is based on the ISI sequence given by

$$ISI_n = t_{n+1} - t_n, n = 1, 2, \cdots, N - 1 \tag{2}$$

First the mean of the ISI sequence is computed via

$$\text{Mean} = (\sum_{n=1}^{N-1} \text{ISI}_n)/(N - 1) \tag{3}$$

Next a new sequence ($L(n)$) is constructed of all the ISIs in the original sequence falling below this mean. We compute the mean (ML) of this new sequence $L(n)$ and define a bursting interval as one falling below ML. To obtain the bursting ratio, we first count the number of ISI below this threshold in the original sequence and then divide it by the total number of ISI's in the original sequence. See Figure 1 for a sample of spike trains with different burst ratios. Notice that trials with burst ratios above 0.5 are bimodally distributed. The burst ratio was computed for each individual spike train and spike trains exceeding a ratio of 0.5 either pre or post stimulus were excluded from analysis (Figure 2).

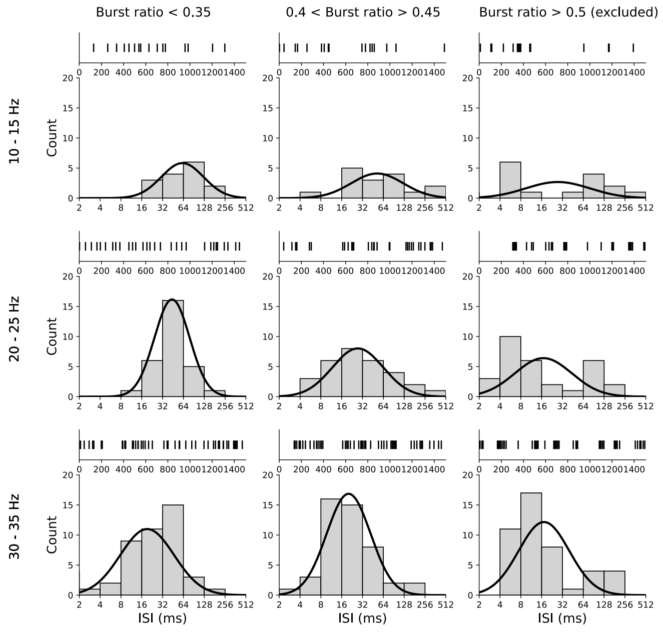


Figure 1: **Bursting ratios across firing rates.**

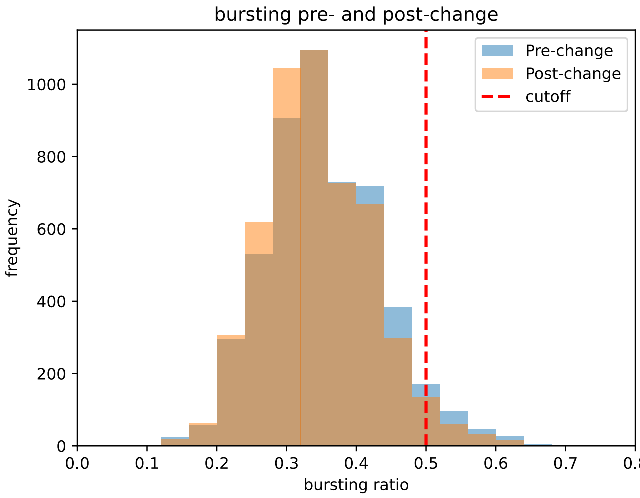


Figure 2: **Bursting before and after stimulus change.**

### 2 Validation

To validate the posterior learned using simulation based inference we performed a validation. First we sampled 100.000 parameters parameter pairs from the prior distribution used during training. For each pair of parameters we obtained an observation by simulating them for 1500ms (to match the pre and post stimulus period length). We then used the posterior to try to infer the original parameters (sampled from the prior). The error was computed as the difference between the recovered and the real parameters. We found that we made larger errors in recovering the parameters from spike trains with fewer spikes (Figure 3a). Moreover, these errors were biased to overestimate the mean and underestimate the variability of the transmembrane current. We thus opted to exclude spike trains containing fewer than 15 spikes (10Hz for 1500ms sampling time). This removed the bias from the inference procedure (Figure 3b).

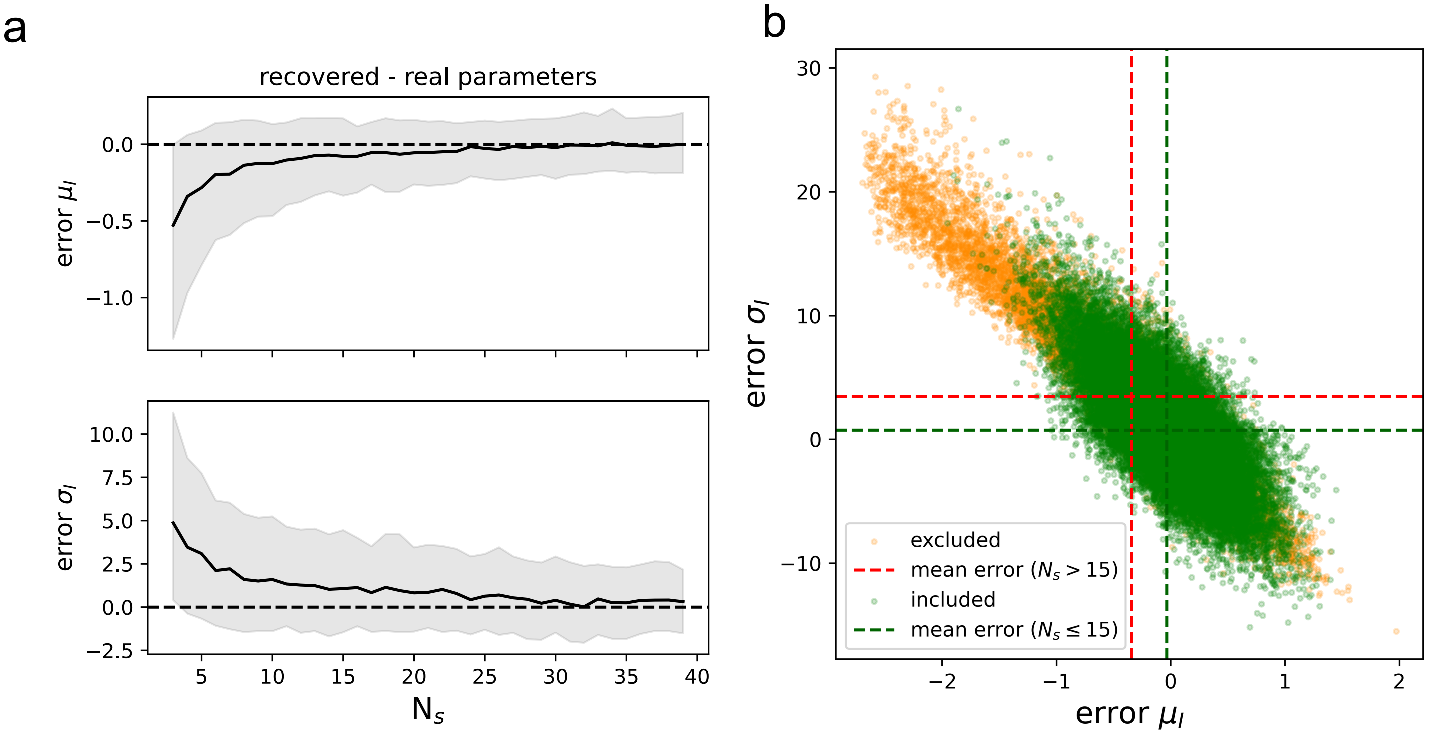


Figure 3: **Validation.** (a) Parameter recovery errors from simulated spike trains containing different numbers of spikes. (b) The recovery bias when including or excluding trials containing more or fewer than 15 spikes.

### 3 Model Specifications

Below are the model specifications according to the framework proposed by Nordlie et al. (2009).

| **A** | **Model Summary** |
| --- | --- |
| **Topology** | - |
| **Connectivity** | - |
| **Populations** | - |
| **Neuron model** | Leaky integrate-and-fire, fixed voltage threshold, fixed absolute refractory time (voltage clamp) |
| **Channel models** | - |
| **Synapse model** | - |
| **Input** | Injected gaussian noise |
| **Measurements** | Spike activity & membrane potential |

| **B** | **Neuron Model** |
| --- | --- |
| **Name** | lif neuron |
| **Type** | Leaky integrate-and-fire |
| **Subthreshold dynamics** | $\tau_\mathrm{m}\frac{dV}{dt}(t) = -(V_\mathrm{m}(t) - E_\mathrm{L}) + RI(t)$ , |
| **Spiking** | if $V_\mathrm{m}(t^*) \geq \Theta \rightarrow V(t^* + \delta t) = V_\mathrm{reset}$<br><br>and emit spike with timestamp $t^*$ |

| **C** | **Neuron Parameters** |
| --- | --- |
| $C_m$ | 1.0 |
| $t_{ref}$ | 0.1 |
| $V_{reset}$ | -70.0 |
| $\tau_m$ | 20.0 |
| $V_{th}$ | -55.0 |
| $E_L$ | -70.0 |

| **D** | **Input** |
| --- | --- |
| **Type** | **Description** |
| Gaussian white noise | $I(t) \sim \mathcal{N}(\mu_\mathrm{I}, \sigma_\mathrm{I})$ |

| **E** | **Measurements** |
| --- | --- |
| **Type** | **Description** |
| Spike times | Time stamps for spiking activity were recorded |
| Membrane potential | Single simulation membrane potential was recorded |
